# Different principles govern different scales of brain folding

**DOI:** 10.1101/851550

**Authors:** Arka N. Mallela, Hansen Deng, Alan Bush, Ezequiel Goldschmidt

**Affiliations:** Department of Neurosurgery, University of Pittsburgh Medical Center, Pittsburgh, Pennsylvania, United States of America; Department of Neurosurgery, Massachusetts General Hospital, Boston, Massachusetts, United States of America

**Keywords:** brain folding, gyrification, fetal MRI, Sylvian Fissure, nonlinear registration

## Abstract

The signature folds of the human brain are formed through a complex and developmentally regulated process. *In vitro* and *in silico* models of this process demonstrate a random pattern of sulci and gyri, unlike the highly ordered and conserved structure seen in the human cortex. Here, we account for the large-scale pattern of cortical folding by combining advanced fetal MRI with nonlinear diffeomorphic registration and volumetric analysis. Our analysis demonstrates that *in utero* brain growth follows a logistic curve, in the absence of an external volume constraint. The Sylvian fissure forms from interlobar folding, where separate lobes overgrow and close an existing subarachnoid space. In contrast, other large sulci, which are the ones represented in existing models, fold through an invagination of a flat surface, a mechanistically different process. Cortical folding is driven by multiple spatially and temporally different mechanisms, therefore regionally distinct biological process may be responsible for the global geometry of the adult brain.

## INTRODUCTION

Mechanisms underlying gyrification of the human cerebral cortex have been investigated for decades. Folding is a dynamic and complex process driven by tangential cortical expansion (Bayly et al. 2014; Kroenke and Bayly 2018) and differential cell growth in and around the subventricular zone (SVZ) (Aydin et al. 2009; Lui et al. 2011; Reillo et al. 2011; Fernández et al. 2016; Borrell 2018). Mechanisms such as a volume constraint from the skull (Welker 1990; Fernández et al. 2016) and tension forces from axons (Van D.C. 1997; Xu et al. 2010) have not received further validation. Tallinen et al. simulated the triple junction folding of the mammalian cortex using a soft elastic bilayers with variable growth rates *in silico* (Tallinen et al. 2014, 2016). The model introduced in this landmark paper produced folds resembling cortical gyri morphologically but did not explain the highly conserved spacing and orientation of gyri. To account for the biology of a preserved cortical pattern, multiple reports have studied folding in ferrets (Xu et al. 2010; Reillo et al. 2011). This is an ideal model to understand the formation of individual gyri and the differences between sulcal and gyral molecular signatures but does not explain the global geometry of the mature telencephalon.

To further guide these models and understand the global pattern of gyrification, analysis of human cortical folding *in vivo* is required. Hill et al. studied postnatal human and macaque brain folding, illustrating that patterned differential expansion in cortex drives an orderly progression of postnatal brain folding(Hill et al. 2010), Garcia et al. studied pre-term and post-natal magnetic resonance imaging (MRI) scans and identified regional differences of cortical growth beginning at an estimated gestational age (GA) of 28 weeks. Notably, increased growth was first observed near the central sulcus from GA 28-30, and subsequently progressing outwards to the frontal and temporal poles (Garcia, Kroenke, et al. 2018). Finally, Rajagopalan et al. utilized *in utero* MRI in a cohort of 40 individuals from GA 20-28 and found similar areas of differential expansion driving cortical folding(Rajagopalan et al. 2011). Here, we build on these analyses by using the latest advancements in fetal MRI reconstruction, fetal atlas construction, and nonlinear diffeomorphic registration.

We hypothesize that cortical folding is driven by multiple spatially and temporally distinct processes. To demonstrate this, we utilized a fetal brain MRI atlas constructed from 81 individuals (Gholipour et al. 2017a) to study changes in the cortical topography in relation to the subarachnoid space at various points of pre-natal maturation, from GA 21 to 38. Our analyses show that interlobar folding, formation of large (named) sulci, and formation of secondary (nonnamed) sulci are morphologically different. Moreover, these proceed in an orderly fashion during development. These findings support the hypothesis that different scales of brain folding are driven by different mechanisms, small and large scale gyrification may obey distinct principles.

## METHODS

### Fetal brain MRI

We utilized a digital fetal brain atlas constructed from 81 healthy fetuses, spanning GA 21 to 38, developed by Gholipour et al. (Gholipour et al. 2017a). Full acquisition details are described in that report. The full content of the atlas can be accessed at http://crl.med.harvard.edu/research/fetal_brain_atlas/.

In brief, the authors acquired structural images using repeated T2-weighted half-Fourier acquisition single shot fast spine echo (T2wSSFSE) series over an acquisition period of 15 to 30 minutes on 3T Siemens Skyra or Trio MRI scanners (Siemens Healthineers, Erlangen, Germany). Images were preprocessed with motion and bias field correction and super-resolution volumetric reconstruction. The atlas was then constructed from volumetric reconstructions utilizing kernel regression over time and symmetric diffeomorphic deformable registration in space. Fetal atlas segmentation of cortical and deep structures was initialized using a previous segmentation developed by Gousias et al. (Gousias et al. 2012, 2013) and propagated to all gestational ages.

### Registration and validation

To identify and quantify week-to-week brain development, we utilized the Symmetric Normalization (SyN) algorithm in the Advanced Normalization Tools (*ANTs*) package to register weekly image pairs (Week 21 to 22, 22 to 23, etc.). This algorithm, a form of symmetric diffeomorphic nonlinear registration, can identify large deformations in tissue morphology while ensuring invertibility, smoothness, and preserving topology (i.e gyri do not intersect one another, brain curvature does not reach an unphysically high value). These registrations are mathematically invertible, allowing for studying week-to-week changes both in the forward and backward directions (Center for History and New Media n.d.; Beg et al. 2005; Avants et al. 2008; Tustison and Avants 2013).

We studied three registration/warping paradigms. For the first two, the registration metric only used image intensity (mutual information) and no additional segmentation or landmark data (that might influence growth predictions). The first registered and warped segmentations from Week 38 to 37, 37 to 36, etc. (“Week 38 to 21”), while the second registered from Week 21 to 22, 22 to 23, etc. (“Week 21 to 38”). For a third method we used both image mutual information and landmark (segmentation) overlap as registration metrics, and registered from Week 38 to 37, 37 to 36 etc. (“Week 38 to 21 with landmarks”)

We used the preexisting segmentations in the Gholipour et al. atlas (Gholipour et al. 2017b) as the ground truth to validate the accuracy of our registration. For the first validation, we warped each preexisting segmentation by 1 week (Week 38 to 37, etc.). For the second, we warped the initial segmentation (Week 38 for the Week 38 to 21 and Week 38 to 21 with landmarks paradigms, Week 21 for the Week 21 to 38 paradigm) through all weeks. We then compared the overlap of the warped segmentations to the preexisting segmentations. using the weighted multiclass Dice similarity coefficient (DSC). The DSC ranges from 0 (no overlap) to 1 (perfect overlap).

As described in the results, the Week 38 to 21 registration paradigm was superior and was used for the remainder of the study. Notably, the only matching criteria used for our final registration was image intensity. No *a priori* segmentation or landmarks were used in our final registration, allowing the algorithm to identify tissue correspondences without outside guidance. Interestingly, when pre-existing segmentations were provided to the registration algorithm, registration accuracy actually worsened (**Tables S1/S2**), suggesting that our registration algorithm may identify correspondences between weeks that do not correspond to conventional boundaries between gyri, sulci, and subcortical structures. We utilized the *ANTsPy* python wrapper around the *ANTs* package and the *Scikit-learn* package in Python (Pedregosa et al. 2011).

### Segmentation

While the original atlas contained several key structures, others, including notable subarachnoid cisterns and sulci, were not included. As such, we algorithmically generated segmentations of the posterior fossa, Sylvian fissure, calcarine sulcus, parietoocipital sulcus and falx cerebri by using the procedure described by Glaister et al. (Glaister et al. 2017). In essence this method identifies all cerebrospinal fluid (CSF) voxels lying between predefined cerebral structures. The posterior fossa space was defined using segmentations of the cerebellum, medulla, pons and midbrain, as well as the associated CSF space. The Sylvian fissure was defined as the CSF space lying between the margins of the frontal operculum, temporal operculum, and supramarginal gyrus. The calcarine sulcus was defined as the sulcus between the cuneus and lingual gyri, and the parietoocipital sulcus as the sulcus between the precuneus and cuneus.

Intracranial volume was defined as all intradural volume. The intracranial space was differentiated into the volumes of the cerebrum, ventricles, and the subarachnoid space. The cerebral volume was defined as the total brain volume lying above the tentorium cerebelli. The brainstem and its associated cisterns were excluded from the analysis since they are not part of the cortical folding process. The ventricles were defined using the preexisting segmentation (Gholipour et al. 2017a). The subarachnoid space was defined as all supratentorial CSF space. This space was further subdivided. The sulcal subarachnoid space was defined as space contained within the sulci/fissures of the brain and which would be directly affected by gyration/sulcation. In contrast, the subdural subarachnoid space, that is, the overlying space between the lateral border of the cerebrum and the dura, reflects how close the brain gets to the skull during development. This was done by constructing a convex hull for each cerebral hemisphere at each gestational point during fetal development. CSF voxels within the shell were termed the sulcal subarachnoid whereas space outside of the shell was subdural subarachnoid (**Figure 1A**)

**Figure 1:**
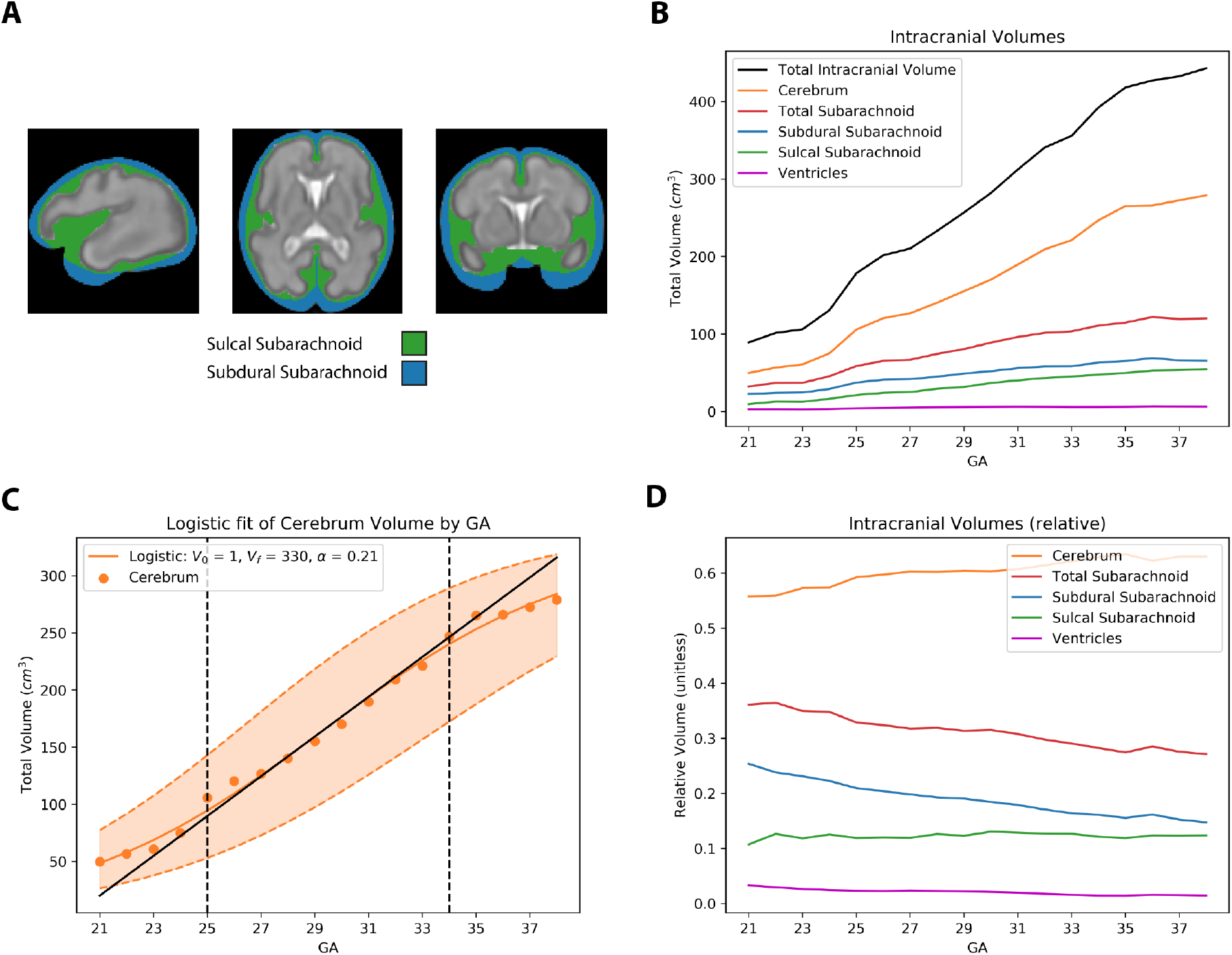
Overall growth of cerebrum and constituent tissues. A) Division of subarachnoid space into sulcal (investing sulci) and subdural (superficial to gyri/sulci). The sulcal subarachnoid is a the space within the sulci/fissures of the brain, while the subdural subarachnoid is the CSF between the outer surface of the brain and inner table of the skull. These spaces were divided by a convex hull around each cerebral hemisphere. B) Total intracranial volume by gestational age. The total volume of the cerebrum increased 6-fold, while the subarachnoid space only grew 4-fold and the ventricles 2-fold. C) Logistic fit for cerebral volume growth. Initial predicted volume (V_0_) was 1 cm^3^, final predicted volume was 330 cm^3^, and logistic growth rate (α) was 0.21, corresponding to a maximum weekly growth rate of 17 cm^3^ per week at approximately 28 weeks. Dashed orange – 1 S.D. error bars, vertical black dashed lines – periodization of brain growth into 3 phases that correspond to folding patterns (see text). Solid orange – logistic fit. Solid black – linear growth in the second phase of growth. D) Relative intracranial volume by gestational age. Cerebral volume expands from 56% to 63%. Note that while there is relative loss in subarachnoid space, this is largely driven, by loss in subdural subarachnoid (25% to 15%), while sulcal subarachnoid is relatively constant at 11-12%. CSF still constitutes approximately 30% of overall cerebral volume at term.

### Volumetric Analysis

We calculated the volumes of total intracranial compartments (intracranial, cerebral, subarachnoid, etc.) and for each analyzed sulcus (Sylvian fissure, parietoocipital sulcus, calcarine sulcus) and the surrounding cortical structures. For global volumetric analysis, we analyzed changes in intracranial compartment volume overtime and performed nonlinear regressions to identify a candidate model for cerebral volume growth. For sulcal volumetric analysis, we calculated the volume of each sulcus and surrounding cortical structures. We analyzed the relationship between sulcus volume and surrounding gyri volumes for each sulci studied.

Neuroimaging data analysis was performed utilizing the *NumPy* and *NiBabel* packages in Python (Python 3.5). Statistical analysis was conducted using the *pandas* package and data plots were generated using *matplotlib* (Matplotlib version 3.1.1) in Python(Oliphant 2007).

### Jacobian Analysis

To map local volumetric changes, the Jacobian determinant was computed at each point for each transformation. The Jacobian determinant quantifies local volumetric change at a given point, with values greater than 1 representing expansion and those less then 1 representing compression. The ANTS algorithm calculates a separate affine transformation and a nonlinear transformation for each registration. The affine transformation accounts for global scaling, transformation, rotation, etc. while the nonlinear component identifies local changes. Here, the Jacobian determinant is calculated from the nonlinear component only. As such, the Jacobian determinant values here is corrected for global volume expansion (scaling). For example, if the registration identified a 10% global expansion, a Jacobian determinant value of 1.1 (+ 10%) represents a total expansion of 20% while a a value of 0.91 (−9%) represents a total expansion of 1%. This was calculated utilizing the *ANTs* toolbox.

### Visualization

We generated 3D surface visualizations to facilitate intuitive understanding of our findings. For figures illustrating sulci and surrounding gyri, we utilized 3D rendering in *itk-snap* (Yushkevich et al. 2006). To illustrate the cortical variation in the Jacobian determinant, we computed a 3D mesh of the cortical surface at each GA using the marching cubes algorithm in the *scikit-image* package and smoothed our mesh using *trimesh*. Jacobian determinant maps were overlaid on this mesh using *nilearn*.

All packages were run on Python 3.7 and utilized under the BSD or GNU license allowing for use in scientific settings.

## RESULTS

### Intracranial expansion by compartment

We segmented each week into various compartments - intracranial volume, total cerebral volume, subarachnoid space, and lateral ventricles. We further divided the subarachnoid space into the subdural subarachnoid space – the space between the lateral border of the cerebrum and dura, reflecting the space between brain and dura, and the sulcal subarachnoid space – the space contained within the sulci/fissures of the brain (**Figure 1A**).

From GA 21 to GA 38 the total volume of the cerebrum increased nearly 6-fold from 50 cm^3^ at GA 21 to 279 cm^3^ at GA 38. Congruently, the total subarachnoid space increased 4-fold from 32 cm^3^ at GA 21 to 120 cm^3^ at GA 38. The ventricles, on the other hand, represented a small volume of 3 cm^3^ at GA 21 grew only 2-fold to 6 cm^3^ throughout gestation. We observed maximum intracranial expansion between weeks 25 and 34. (**Figure 1B)**.

We found that a logistic growth model best fits the cerebral volume over gestation **(Figure 1C)**. This parcellates growth into 3 phases. From Week 21 to 25 growth is relatively slow, but the growth rate increases between week 25 and 34, with a final tapering. The model predicts a final (term) volume of 330 cm^3^ and a maximum weekly growth rate of 17 cm^3^ per week at approximately 28 weeks.

In relative terms, cerebral volume expands from 56% to 63% of intracranial volume. While total subarachnoid relative volume decreases from 36% to 27%, this is largely driven by loss in subdural subarachnoid space (25% to 15%). Importantly, almost a third of the intracranial volume corresponds to subdural space at the end of gestation. Thus, cortical folding of the brain occurs in the presence of abundant subdural subarachnoid space and a lack of an external volume constraint. Interestingly the sulcal subarachnoid relative volume remains essentially constant from 11% to 12%, suggesting that globally the folding process creates and obliterates equal relative amounts of subarachnoid space. (**Figure 1D**).

### The Sylvian fissure folds differently than other sulci

The majority of cerebral sulci are elongated depressions between two raised gyri. This type of folding occurs in parallel with intracranial expansion and is the most prevalent type of sulcal formation. As the bordering gyri increase in volume, the sulcus deepens, and both should show a correlated increase in total volume. To analyze this form of folding, we segmented each sulcus/fissure and identified the surrounding gyri.

We used the left parieto-occipital and calcarine sulci to characterize this form of folding. In these sulci, both the volumes of the sulcal edges and the sulcal subarachnoid space increased with time, which resulted in a direct correlation (r = 0.99 and r = 0.95, respectively; **Figure 2 center, right**).

**Figure 2:**
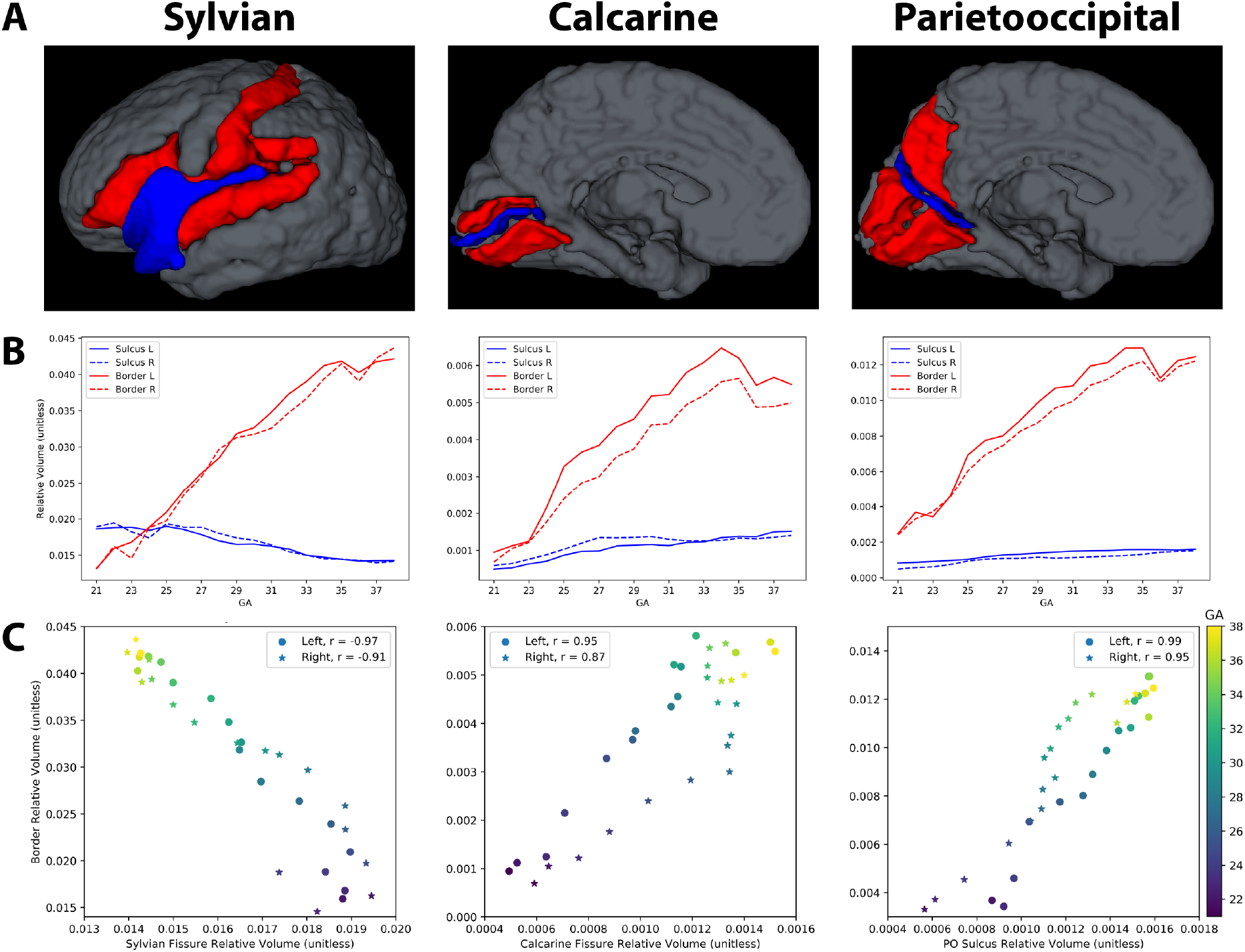
Sylvian fissure develops differently than other fissures. A) Illustrations of sulci (blue) and bordering gyri (red) at Week 33. B) Relative volume of sulcus compared to that of bordering gyri. For the calcarine and parietooccipital sulci, both sulcus and gyri increase in relative volume, unlike in the Sylvian fissure. C) Correlation between sulcus relative volume and bordering gyri relative volume. The Sylvian fissure (r = − 0.97) demonstrates a strong negative correlation while the parietooccipital (r = 0.99) and calcarine (r = 0.95). demonstrate strong positive correlations. The Sylvian fissures shrinks as the opercula grow while the other sulci grow with their bounding gyri.

The Sylvian fissure, on the other hand, appears to form from a flat depression that is closed through the convergence of two edges. It first appears on the lateral aspect of the cerebral hemisphere before other folds are present. Subsequently, there is a gradual confluence of the anterior and posterior edges of the depression which closes the depression. These edges develop into the frontal and temporal opercula and the depression itself forms the insula (**Figure 3**). This mechanism of folding is completely different than that of other sulci and fissures, exemplified here by the parietooccipital and calcarine fissures. The growth of these Sylvian opercula is inversely correlated with the growth of the Sylvian subarachnoid space (r = −0.97; **Figure 2, left**). The Sylvian fissure forms through a convergent closure as opposed to the invagination characteristic of other sulci – as the bordering gyri grow, the Sylvian fissure shrinks.

**Figure 3:**
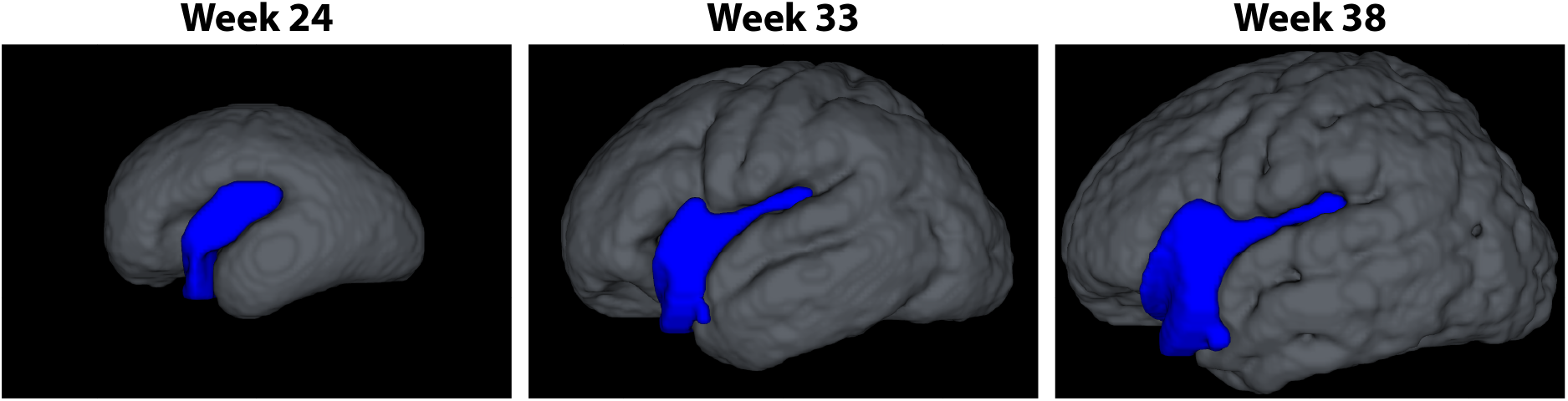
Sylvian fissure closes over gestation. The Sylvian fissure begins as a flat depression and closes over time through the overgrowth of the opercular gyri. This process is largely complete by Week 33 or earlier. This stands in contrast to the other sulci of the brain and is the prime example of interlobar folding

### Spatiotemporally distinct processes fold the brain at multiple scales

Relative volumetric change (as determined by the Jacobian determinant) were quantified in both the left and right hemispheres and are rendered in **Figure 4**. As described in the methods, the Jacobian is corrected for *global* volume growth, so local loss in volume still may represent overall growth.

**Figure 4:**
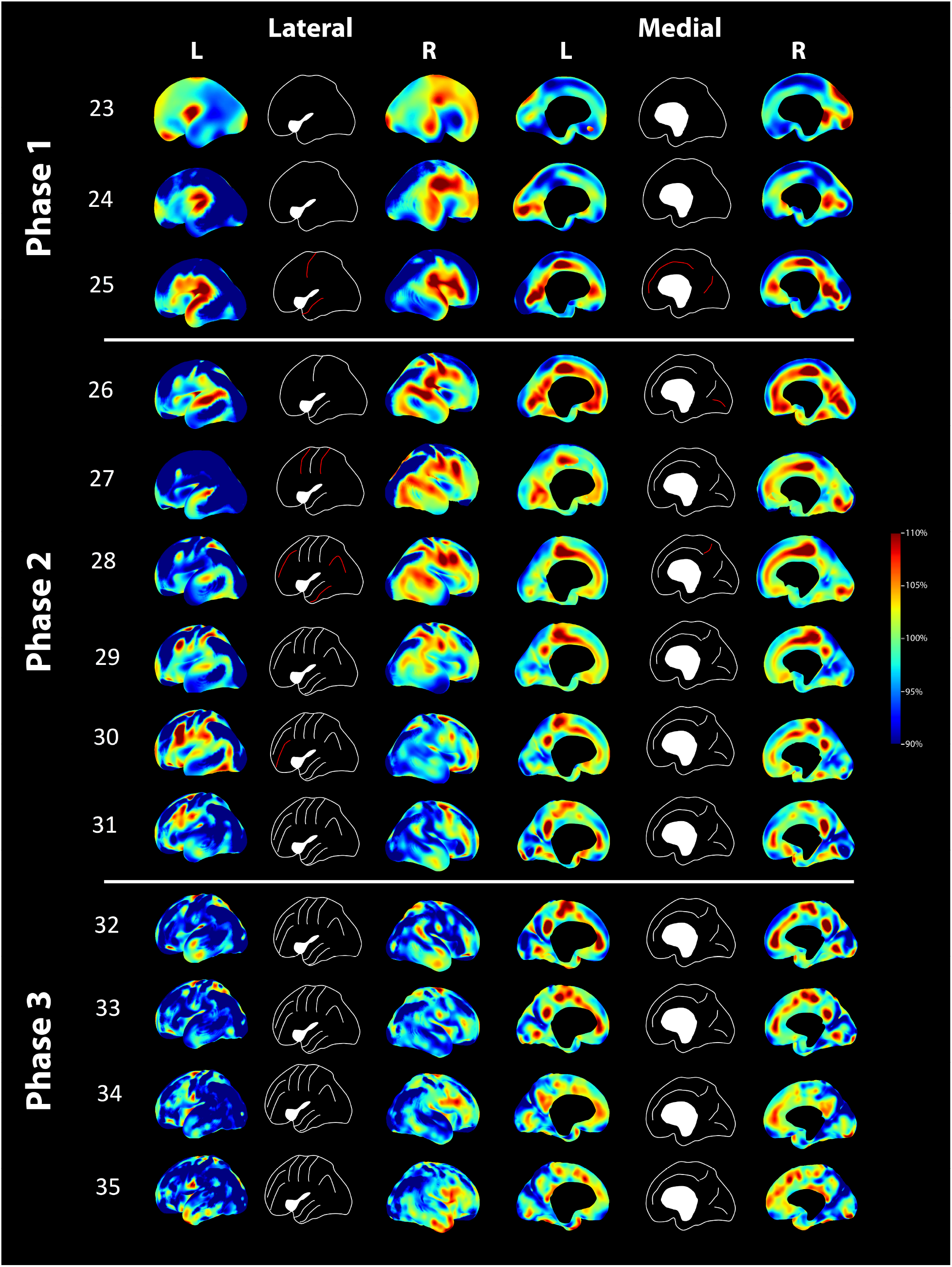
Jacobian analysis (week-to-week relative volume expansion, correcting for overall volume growth). *Left:* lateral cerebral surface. *Right:* medial cerebral surface. *Rows* – gestational age. The central column in each group demonstrates existing (white) or new (red) sulci at each week. In phase 1, folding is primarily interlobar with the majority of folding occurring in the perisylvian areas (frontal, parietal, and temporal opercula), forming and closing the Sylvian fissure. In doing so, the major lobes are demarcated In phase 2, folding forms the major named sulci of the brain, proceeding in an orderly fashion from the central sulcus outward. On the medial surface, the parietooccipital and calcarine sulci finish folding and the cingulate sulcus elongates and serves as a source for additional volume expansion. By phase 3 (around GA week 32), the major sulci have completed forming and smaller, non-named sulci are folding. The only area of persistent area growth by week 35 onwards (weeks 36-38 not pictured) is the temporal pole.

#### Lateral hemispheric surface

Initially, the primary area of folding is the frontal, parietal, and temporal opercula, with closure of the Sylvian fissure. From GA 23 to 25, the depression that becomes the future insula becomes closed off in the posterior aspect, and the gross morphology of the major lobes is formed. Volumetric change was first observed at the supramarginal gyrus starting in GA 23, then moved anteriorly to the frontal lobe and laterally in the direction of the temporal pole. This “C” shaped expansion originates from the area that will become the supramarginal gyrus eventually reaching the future pre-central gyrus. From GA 25 to 31, focal volume growth was observed around the forming Sylvian fissure and the incipient central sulcus. In the frontal and parietal lobes, the central sulcus forms first and then sulcation proceeds in an orderly fashion outward from the central sulcus. The central sulcus is apparent by GA 26, the pre and post central sulci by GA 28-29, the intraparietal sulcus by GA 29, and the superior and inferior frontal sulci by GA 31. In the temporal lobe, the superior temporal sulcus appears to form first, by approximately GA 29 and then inferior temporal sulcus appears to be forming at GA 31. At GA 32, the major sulci of the brain have formed and are clearly recognizable. However, folding continues to proceed with the formation of additional convolutions, but there does not appear to be an orderly progression of folding, with only the temporal pole and frontal operculum appearing to have consistent volume expansion.

#### Medial hemispheric surface

Spatiotemporal mapping of the medial hemispheres revealed that the cingulum underwent the greatest differential volumetric change. By GA 24, a yet unfolded brain exhibited increased growth in 3 regions involving the medial frontal lobe, the center of the cingulum, and the occipital lobe in what will become the anatomical parieto-occipital sulcus. By week 26, these high-growth areas formed an inverted “U,” marked posteriorly by a low-growth area in the precuneus. Similar to development observed on the lateral bi-hemispheric surfaces, there was volumetric expansion of the paracentral lobule (the representation of the pre and post central gyri on the medial surface) slowing by week 31-33. By Week 29, a clear parieto-occipital sulcus is visible. Interestingly volumetric growth around the cingulate sulcus is the norm, only slowing near the end. At a later stage, expansion is constrained to a few dispersed areas, without a clear pattern in folding, similar to the growth on the lateral hemisphere.

#### Ventricular zone

**Figure 4** demonstrates significant growth at the cortical surface. Interestingly, while the subplate zone also demonstrates volumetric expansion, the ventricular zone (VZ) does not (**Figure S1**). There is a clear boundary running through the SVZ dividing this area of relative expansion vs. relative contraction in the VZ. This dichotomy between superficial and deep structure persists across all GA in this study.

### Validation of registration

In order to validate our registration method, we compared three registration/warp paradigms – Week 38 to 21 (with intensity information only), Week 21 to 38 (with intensity information only), and Week 38 to 21 with landmarks. We warped segmentations week-to-week (**Table S1**) and cumulative across all weeks starting from either Week 21 or 38 (first/last weeks) (**Table S2**) and compared them to the preexisting segmentation using the weighted multiclass Dice similarity coefficient (range: 0 – no overlap, to 1 – perfect overlap).

For the registration used for our final experiments (Week 38 to 21), the week-to-week comparison Dice coefficient ranged from 0.89 to 0.94 (**Table S1**), while the cumulative comparison – warping the Week 38 segmentation across all weeks – ranged from 0.9 (week 38 to 37) to 0.72 (week 38 warped via concatenated transformations to week 21) (**Table S2**).

Overlap when segmentations were warped in the forward direction, from week 21 to 38, were comparable in the week-to-week comparison, ranging from 0.89 to 0.94 (**Table S1**). However, in the cumulative comparison, this paradigm fared slightly worse, with Dice coefficients ranging from 0.9 (week 21 to 22) to 0.68 (week 21 warped via concatenated transformations), reflecting greater accumulated registration error in this paradigm (**Table S2**).

Interestingly, when the registration algorithm was given the preexisting segmentation as a landmark in addition to image intensity, overlap worsened both in week-to week (0.87 – 0.91, **Table S1**) and cumulative comparisons (0.9 to 0.56, **Table S2**).

## DISCUSSION

In the second and third trimesters of gestation, the cerebrum undergoes regulated expansion and the formation of gyri and sulci. Previous investigations have proposed multiple mechanisms for the folding of a smooth cortical plate into a highly convoluted cortex (Van D.C. 1997; Hilgetag and Barbas 2006; Herculano-Houzel et al. 2010; Reillo et al. 2011; Zilles et al. 2013; Dubois et al. 2014; Tallinen et al. 2014, 2016; Tallinen and Biggins 2015; Fernández et al. 2016; Borrell 2018; Garcia, Kroenke, et al. 2018). Most of these models have rendered a prototypical sulcus, but few have studied the development of the entire human cortex *in vivo* (Hill et al. 2010; Rajagopalan et al. 2011, 2012; Garcia, Kroenke, et al. 2018). None have systematically mapped the spatiotemporal folding of the cortex in the second and third trimesters. Our present study does so, specifically highlighting on a week-to-week basis how distinct spatiotemporal processes gyrify the brain *in utero*.

We elected to analyze the Gholipour atlas (Gholipour et al. 2017a) as it was the highest resolution available dataset and provided much higher signal-to-noise ratio than an individual fetal MRI. Given that this atlas was constructed as a normative group mean of 81 fetuses, we expect that our findings will translate to the individual setting.

### Cerebral growth is internally constrained

Our study demonstrated that both the intra-axial and extra-axial spaces grow significantly during gestation (**Figure 1**). At birth, more than 15% of the total intracranial volume corresponds to subdural subarachnoid space, indicating a significant layer of CSF between the cerebrum and inner table of the skull. This further strongly suggests that cortical folding of the brain is not driven by a mechanical extra-axial constraint. Moreover, there is increasing proteomic evidence that the CSF provides a proliferative niche during embryonic development via fibroblast growth factor (Martín et al. 2006) and insulin-like growth factor (Lehtinen et al. 2011), among other signaling molecules.

Yet, our analysis indicates that cerebral growth still slows by late gestation, following a logistic curve, with a term volume of 330 cm^3^, similar to previous reports in the literature(Holland et al. 2014). Similarly, Dubois et al. found that perinatal brain growth follows a Gompertzian curve, an asymmetric relative of the logistic curve (Dubois et al. 2019). Both models indicate a constraint to unrestrained growth – such as cell-to-cell signaling, internal mechanical constraints, among other mechanisms(Laird 1965; West and Newton 2019). Despite a lack of an *external* volumetric constraint, other internal phenomena alter and shape the growth of the cerebrum.

### Different principles govern different scales of brain folding

The development of the Sylvian fissure, the exemplar of interlobar folding and the main depression responsible for the global configuration of the mature brain, is a fundamentally different process than large sulci formation. Morphologically, the Sylvian fissure *closes* while the major other sulci *deepen*. This is apparent both qualitatively from the maps of volume expansion (**Figure 4**) and quantitatively (**Figure 2**). The borders of the calcarine and parietoocipital sulci expand outwards while the sulci deepen – both expand in volume in a correlated fashion. In contrast, as the opercular gyri grow, the Sylvian fissure shrinks – in an anticorrelated fashion (**Figures 2 and 3**). This suggests that different biophysical, genetic, and cellular growth mechanisms fold the Sylvian fissure.

The presence of a distinct mechanism for interlobar folding is supported by two observations. First, global malformations of cortical development can have sulci without the Sylvian fissure or vice versa. For example, in alobar holoprosencephaly, sulci can be present, but there is no global structure to the brain, let alone a Sylvian fissure. In contrast, in lissencephaly, there is a Sylvian fissure but few to no other sulci. Finally, even in polymicrogyria, often a Sylvian fissure is present even though all other sulci are malformed (Sarnat and Flores-Sarnat 2016). This suggests that the mechanisms for interlobar folding and sulcation are different. Second, recent work by Bush et al. (Bush et al. 2019), seeks to explain why no middle cerebral artery branches cross the Sylvian fissure, when *every other sulcus* has traversing arteries. The authors developed a novel convergence index calculated as the ratio of shortest cortical path to physical distance. Using this they demonstrate that the frontal and temporal opercula have the highest degree of convergence of any cortical structure, implying that they begin in fetal life distant from one another and the converge over time. In contrast, the borders of other sulci do not converge to such a high degree. This again suggests that interlobar folding and large sulcation are distinct processes.

Following interlobar folding, the brain appears to undergo an orderly progression of folding during large sulcation, beginning with the formation of the central sulcus and superior temporal sulcus, and progressing outwards/inferiorly. This is driven by regions of higher expansion compared to the surrounding cortex. Garcia et al. described such regions of high growth being located around the insula and migrating outward from the central sulcus (Garcia, Robinson, et al. 2018). Likewise, both Rajagopalan et al. (Rajagopalan et al. 2011) and Dubois et al. (Dubois et al. 2019) found a similar ordered pattern of folding during the late second and early third trimesters, while Garcia et al. demonstrated this outward progression from the central sulcus in preterm but postnatal babies. Interestingly, they analyzed changes in curvature or spatial frequencies of gyri, instead of volumetric changes analyzed here. All three approaches reached similar conclusions despite different methods of analysis, lending further credence to this hypothesis. A purely mechanical explanation cannot account for the highly conserved and ordered nature of this process.

In contrast, the formation of secondary, non-named sulci, such as branches off the superior/inferior frontal sulci and the variable branching of the intraparietal sulcus form later in neurodevelopment. Following GA week 32 we did not observe discrete areas of consistent relative volumetric expansion (**Figure 4**). These sulci are less conserved than the large sulci, exhibiting greater variability between individuals (Thompson et al. 1996; Caspers et al. 2006). This form of folding may be the result of tangential expansion and mechanical creasing, as proposed by Tallinen et al. (Tallinen et al. 2014, 2016; Tallinen and Biggins 2015). Thus, we find that patterns of large-scale folding are qualitatively different from that of secondary sulcation.

Our analysis suggests that the 3 phases of the logistic curve may correspond to the three types of folding – interlobar folding (exemplified by the formation of the Sylvian fissure) until approximately GA 25 weeks (**Figure 4 – Phase 1**), large sulcation (e.g. the central sulcus, pre/post central sulci, temporal sulci) from GA 26 to 31 weeks (**Figure 4 – Phase 2**), and secondary sulcation (**Figure 4 – Phase 3**), from 32 weeks to term. Others have reported a similar periodization, although nomenclature varies, particularly as to what constitutes primary and secondary sulcation. (Chi et al. 1977; Rajagopalan et al. 2011; Clouchoux et al. 2012; Habas et al. 2012; Wright et al. 2014).

### Asymmetries and divisions in growth

We additionally noted a left/right asymmetry in our Jacobian analysis. A left-right asymmetry was first noticed by direct observation of 507 fetal brains by Chi et al., who noticed that the superior frontal and temporal gyri appeared one or two weeks before on the right side (Chi et al. 1977). These findings were later confirmed by Leroy et al., who also described lateral asymmetries specific to humans and that persist during and after gestation (Leroy et al. 2015). This may speak to the genetic nature of the folding process.

Finally, we note that majority of volumetric expansion occurs in the cortex and subplate, with comparatively little to no expansion occurring in the ventricular zone from GA weeks 21 to 38 (**Figure S1**). There appears to be a boundary running through the subventricular zone that divides areas of relative expansion and lack of expansion. Our atlas does not delineate the SVZ exactly or the boundary between the outer and inner SVZ, so we are unable to assess exactly whether our boundary corresponds to a cellular boundary. Nevertheless, this line dividing areas of expansion and lack of expansion is consistent with both a mechanical bilayer tangential growth explanation and an outer SVZ cell growth model but is less consistent with axonal tension models.

### Underlying patterning is required to explain phases of folding

A purely mechanical explanation is insufficient to explain our and others’ findings regarding interlobar folding and large sulcation but may explain secondary sulcation. Findings from other developmental studies suggest that this may be driven by underlying genetic or proteomic patterning in the ventricular zone in the first and second trimesters (Walsh 1999; Lohmann et al. 2008; Reillo et al. 2011; Elsen et al. 2013; de Juan Romero et al. 2015). Nonuniform expression of transcription or growth factors in the ventricular zone may unlock greater growth at a later phase of development, once intermediate progenitor cells have migrated to the subventricular zone. Cells found in of high growth in the Jacobian analysis may have unique expression profiles and are prime targets for tissue analysis.

Mechanical phenomena can explain *how* sulci form, but not where and when they form. Further, less conserved secondary folding may be a purely mechanical phenomenon.

## CONCLUSIONS

Brain folding is not a homogenous process and different principles are required to explain the different scales of folding. Using *in utero* fetal MRI, we demonstrate that interlobar folding, large sulcation, and secondary sulcation are morphologically different. A purely mechanical explanation cannot explain the highly patterned nature of these processes.

## Acknowledgements

We thank the authors of Gholipour et al. (Gholipour et al. 2017a) for making their fetal atlas database publicly available to the scientific community and supporting work such as this.

## Data availability

The fetal brain atlas developed by Gholipour et al. (Gholipour et al. 2017a). can be accessed at http://crl.med.harvard.edu/research/fetal_brain_atlas/. All code utilized for this report will be made freely available upon request to the authors.

## Supplementary Figures

**Figure S1:**
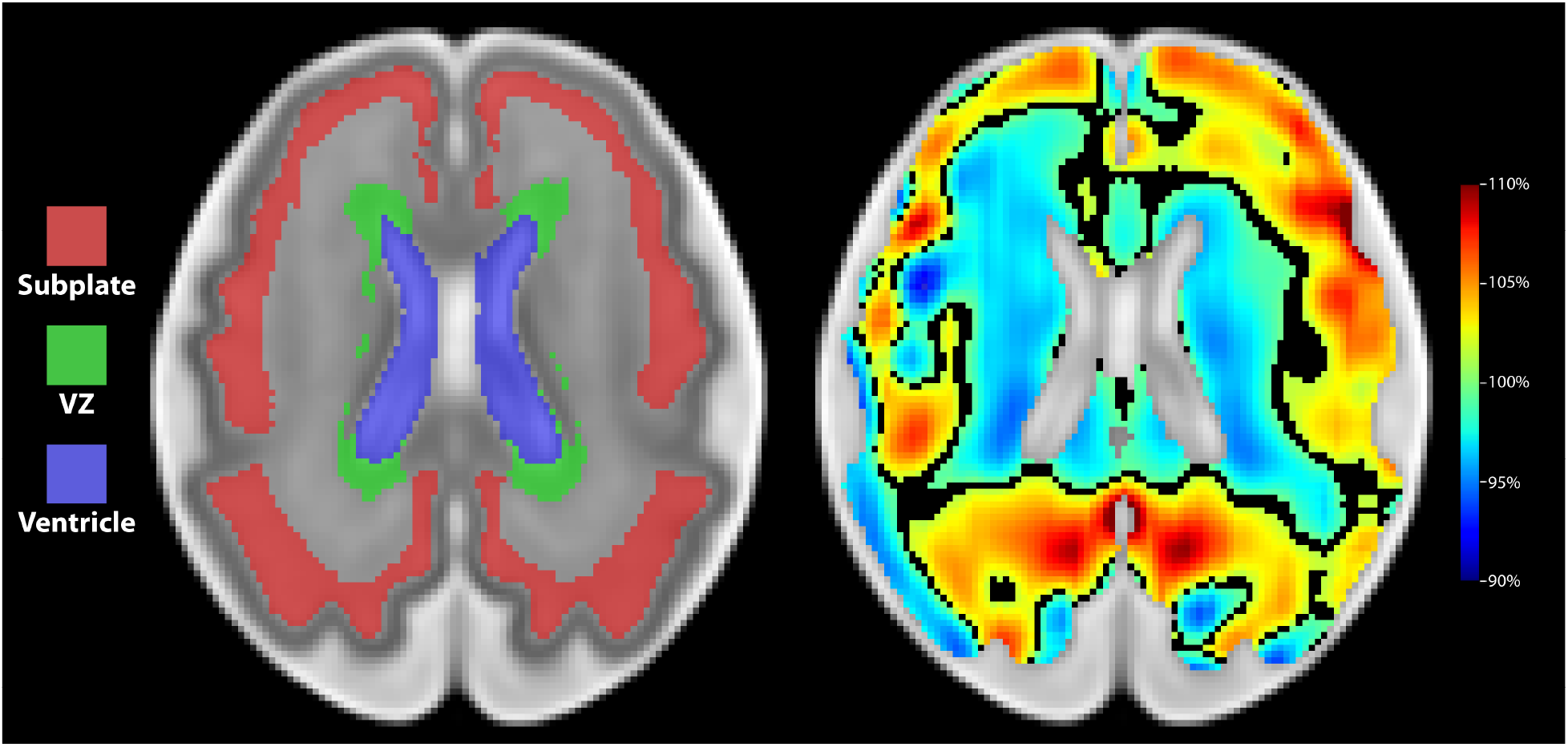
Volume expansion is at the cortical plate and subplate. Axial slice for Week 30 at the level of the lateral ventricles. **Left –** segmentations of the subplate, ventricular zone (VZ), and lateral ventricles. **Right –** Jacobian determinant image with scale bar to right. Black lines represent areas of zero nonlinear growth (dividing areas of relative expansion and lack of expansion). Note the area of expansion largely constrained to the cortical and subplate. The ventricular zone is largely quiescent.

**Table S1.** Registration validation: Comparing segmentation overlap (as measured by Dice similarity coefficient) when warped by one week between different registration/warp paradigms. We performed three registration paradigms for the nonlinear diffeomorphic registration as described in the text. In the first, using image intensity only (image mutual information), we registered from Week 38 to 37, 37 to 36, and so on (“Week 38 to 21”). Second, also using image intensity only), we registered from Week 21 to 22, 22 to 23, and so on (“Week 21 to 38”). Finally we performed the first registration again (Week 38 to 21) but provided the registration algorithm with landmarks from the preexisting segmentations from the Gholipour et al. atlas (“Week 38 to 21 with landmarks”). Each week had preexisting segmentations from the Gholipour et al. atlas. We used these as the ground truth and compared them to warped segmentation. In Table S1, we warped each segmentation by 1 week using the paradigms described above (i.e Week 38 segmentation warped to Week 37, 37 to 36, etc.) We then computed the overlap of the segmentations using the weighted multiclass Dice similarity. The first two paradigms that were only intensity-based (Week 38 to 21 and Week 21 to 38) were largely comparable in overlaps, while Week 38 to 21 with landmarks appeared to have slightly poorer registration accuracy. The results in this paper are based off the Week 38 to 21 registration paradigm.

| GA | Week 38 to 21 | Week 21 to 38 | Week 38 to 21 with landmarks |
| --- | --- | --- | --- |
| 21 | 0.90 | 1.00 | 0.88 |
| 22 | 0.90 | 0.90 | 0.89 |
| 23 | 0.91 | 0.90 | 0.80 |
| 24 | 0.91 | 0.90 | 0.86 |
| 25 | 0.92 | 0.91 | 0.90 |
| 26 | 0.93 | 0.92 | 0.91 |
| 27 | 0.93 | 0.93 | 0.91 |
| 28 | 0.94 | 0.93 | 0.91 |
| 29 | 0.93 | 0.93 | 0.90 |
| 30 | 0.93 | 0.93 | 0.87 |
| 31 | 0.94 | 0.93 | 0.89 |
| 32 | 0.94 | 0.93 | 0.91 |
| 33 | 0.94 | 0.94 | 0.88 |
| 34 | 0.94 | 0.94 | 0.88 |
| 35 | 0.89 | 0.94 | 0.88 |
| 36 | 0.90 | 0.89 | 0.90 |
| 37 | 0.90 | 0.90 | 0.89 |
| 38 | 1.00 | 0.90 | 1.00 |

**Table S2.** Registration validation: Comparing segmentation overlap (as measured by Dice similarity coefficient) when warped over all weeks between different registration/warp paradigms. We performed three registration paradigms and validated them with the segmentation overlap, as measured by the Dice similarity coefficient. In table S2, we warped the initial segmentation from the atlas (Week 38 for the Week 38 to 21 and Week 38 to 21 with landmarks paradigms, Week 21 for the Week 21 to 38 paradigm) through all weeks. For example, we warped the week 38 segmentation to week 37, then that warped segmentation to week 36, etc. We then computed the overlap of the segmentations using the weighted multiclass Dice similarity. Lower, though still acceptable, overlap scores were seen the further the initial segmentation was warped, suggesting an accumulation of error. The final accumulated error was slightly lower in the Week 38 to 21 paradigm than the Week 21 to 38 paradigm. Interestingly, the Week 38 to 21 with landmarks paradigm had significantly worse registration accuracy when compared to the intensity-only paradigms, suggesting that providing the preexisting segmentations directly to the registration algorithm worsened its performance. Given these results, we used the Week 38 to 21 paradigm for the remainder of the paper.

| GA | Week 38 to 21 | Week 21 to 38 | Week 38 to 21 with landmarks |
| --- | --- | --- | --- |
| 21 | 0.72 | 1.00 | 0.56 |
| 22 | 0.72 | 0.90 | 0.59 |
| 23 | 0.73 | 0.88 | 0.60 |
| 24 | 0.73 | 0.86 | 0.63 |
| 25 | 0.74 | 0.85 | 0.67 |
| 26 | 0.75 | 0.83 | 0.69 |
| 27 | 0.77 | 0.82 | 0.71 |
| 28 | 0.78 | 0.80 | 0.72 |
| 29 | 0.80 | 0.78 | 0.75 |
| 30 | 0.80 | 0.78 | 0.77 |
| 31 | 0.82 | 0.76 | 0.78 |
| 32 | 0.83 | 0.75 | 0.80 |
| 33 | 0.84 | 0.73 | 0.81 |
| 34 | 0.86 | 0.72 | 0.84 |
| 35 | 0.87 | 0.71 | 0.87 |
| 36 | 0.88 | 0.70 | 0.88 |
| 37 | 0.90 | 0.70 | 0.90 |
| 38 | 1.00 | 0.68 | 1.00 |

## References

Avants BB, Epstein CL, Grossman M, Gee JC. 2008. Symmetric diffeomorphic image registration with cross-correlation: evaluating automated labeling of elderly and neurodegenerative brain. Med Image Anal. 12:26–41.

Aydin S, Yilmazlar S, Aker S, Korfali E. 2009. Anatomy of the floor of the third ventricle in relation to endoscopic ventriculostomy. Clin Anat. 22:916–924.

Bayly P V., Taber LA, Kroenke CD. 2014. Mechanical forces in cerebral cortical folding: A review of measurements and models. J Mech Behav Biomed Mater. 29:568–581.

Beg MF, Miller MI, Trouvé A, Younes L. 2005. Computing large deformation metric mappings via geodesic flows of diffeomorphisms. Int J Comput Vis. 61:139–157.

Borrell V. 2018. How cells fold the cerebral cortex. J Neurosci. 38:776–783.

Bush A, Nuñez M, Brisbin AK, Friedlander RM, Goldschmidt E. 2019. Spatial convergence of distant cortical regions during folding explains why arteries do not cross the sylvian fissure. J Neurosurg JNS. 1–10.

Caspers S, Geyer S, Schleicher A, Mohlberg H, Amunts K, Zilles K. 2006. The human inferior parietal cortex: Cytoarchitectonic parcellation and interindividual variability. Neuroimage. 33:430–448.

Center for History and New Media. n.d. Zotero Quick Start Guide [WWW Document]. URL http://zotero.org/support/quick_start_guide

Chi JG, Dooling EC, Gilles FH. 1977. Gyral development of the human brain. Ann Neurol. 1:86–93.

Clouchoux C, Kudelski D, Gholipour A, Warfield SK, Viseur S, Bouyssi-Kobar M, Mari JL, Evans AC, Du Plessis AJ, Limperopoulos C. 2012. Quantitative in vivo MRI measurement of cortical development in the fetus. Brain Struct Funct. 217:127–139.

de Juan Romero C, Bruder C, Tomasello U, Sanz-Anquela JM, Borrell V. 2015. Discrete domains of gene expression in germinal layers distinguish the development of gyrencephaly. EMBO J. 34:1859–1874.

Dubois J, Dehaene-Lambertz G, Kulikova S, Poupon C, Hüppi PS, Hertz-Pannier L. 2014. The early development of brain white matter: A review of imaging studies in fetuses, newborns and infants. Neuroscience.

Dubois J, Lefèvre J, Angleys H, Leroy F, Fischer C, Lebenberg J, Dehaene-Lambertz G, Borradori-Tolsa C, Lazeyras F, Hertz-Pannier L, Mangin JF, Hüppi PS, Germanaud D. 2019. The dynamics of cortical folding waves and prematurity-related deviations revealed by spatial and spectral analysis of gyrification. Neuroimage. 185:934–946.

Elsen GE, Hodge RD, Bedogni F, Daza RAM, Nelson BR, Shiba N, Reiner SL, Hevner RF. 2013. The protomap is propagated to cortical plate neurons through an Eomes-dependent intermediate map. Proc Natl Acad Sci U S A. 110:4081–4086.

Fernández V, Llinares-Benadero C, Borrell V. 2016. Cerebral cortex expansion and folding: what have we learned? EMBO J. 35:1021–1044.

Garcia KE, Kroenke CD, Bayly P V. 2018. Mechanics of cortical folding: Stress, growth and stability. Philos Trans R Soc B Biol Sci.

Garcia KE, Robinson EC, Alexopoulos D, Dierker DL, Glasser MF, Coalson TS, Ortinau CM, Rueckert D, Taber LA, Van Essen DC, Rogers CE, Smysere CD, Bayly P V. 2018. Dynamic patterns of cortical expansion during folding of the preterm human brain. Proc Natl Acad Sci U S A. 115:3156–3161.

Gholipour A, Rollins CK, Velasco-Annis C, Ouaalam A, Akhondi-Asl A, Afacan O, Ortinau CM, Clancy S, Limperopoulos C, Yang E, Estroff JA, Warfield SK. 2017a. A normative spatiotemporal MRI atlas of the fetal brain for automatic segmentation and analysis of early brain growth. Sci Rep. 7:476.

Gholipour A, Rollins CK, Velasco-Annis C, Ouaalam A, Akhondi-Asl A, Afacan O, Ortinau CM, Clancy S, Limperopoulos C, Yang E, Estroff JA, Warfield SK. 2017b. A normative spatiotemporal MRI atlas of the fetal brain for automatic segmentation and analysis of early brain growth. Sci Rep. 7.

Glaister J, Carass A, Pham DL, Butman JA, Prince JL. 2017. Automatic falx cerebri and tentorium cerebelli segmentation from Magnetic Resonance Images. Proc SPIE--the Int Soc Opt Eng. 10137.

Gousias IS, Edwards AD, Rutherford MA, Counsell SJ, Hajnal J V, Rueckert D, Hammers A. 2012. Magnetic resonance imaging of the newborn brain: manual segmentation of labelled atlases in term-born and preterm infants. Neuroimage. 62:1499–1509.

Gousias IS, Hammers A, Counsell SJ, Srinivasan L, Rutherford MA, Heckemann RA, Hajnal J V., Rueckert D, Edwards AD. 2013. Magnetic Resonance Imaging of the Newborn Brain: Automatic Segmentation of Brain Images into 50 Anatomical Regions. PLoS One. 8.

Habas PA, Scott JA, Roosta A, Rajagopalan V, Kim K, Rousseau F, Barkovich AJ, Glenn OA, Studholme C. 2012. Early folding patterns and asymmetries of the normal human brain detected from in utero MRI. Cereb Cortex. 22:13–25.

Herculano-Houzel S, Mota B, Wong P, Kaas JH. 2010. Connectivity-driven white matter scaling and folding in primate cerebral cortex. Proc Natl Acad Sci U S A. 107:19008–19013.

Hilgetag CC, Barbas H. 2006. Role of mechanical factors in the morphology of the primate cerebral cortex. PLoS Comput Biol. 2:146–159.

Hill J, Inder T, Neil J, Dierker D, Harwell J, Van Essen D. 2010. Similar patterns of cortical expansion during human development and evolution. Proc Natl Acad Sci U S A. 107:13135–13140.

Holland D, Chang L, Ernst TM, Curran M, Buchthal SD, Alicata D, Skranes J, Johansen H, Hernandez A, Yamakawa R, Kuperman JM, Dale AM. 2014. Structural growth trajectories and rates of change in the first 3 months of infant brain development. JAMA Neurol. 71:1266–1274.

Kroenke CD, Bayly P V. 2018. How forces fold the cerebral cortex. J Neurosci. 38:767–775.

Laird AK. 1965. Dynamics of tumour growth: Comparison of growth rates anid extrapolation of growth curve to one cell. Br J Cancer. 19:278–291.

Lehtinen MK, Zappaterra MW, Chen X, Yang YJ, Hill AD, Lun M, Maynard T, Gonzalez D, Kim S, Ye P, D’Ercole AJ, Wong ET, LaMantia AS, Walsh CA. 2011. The cerebrospinal fluid provides a proliferative niche for neural progenitor cells. Neuron. 69:893–905.

Leroy F, Cai Q, Bogart SL, Dubois J, Coulon O, Monzalvo K, Fischer C, Glasel H, Van Der Haegen L, Bénézit A, Lin CP, Kennedy DN, Ihara AS, Hertz-Pannier L, Moutard ML, Poupon C, Brysbaert M, Roberts N, Hopkins WD, Mangin JF, Dehaene-Lambertz G. 2015. New human-specific brain landmark: The depth asymmetry of superior temporal sulcus. Proc Natl Acad Sci U S A. 112:1208–1213.

Lohmann G, Von Cramon DY, Colchester ACF. 2008. Deep sulcal landmarks provide an organizing framework for human cortical folding. Cereb Cortex. 18:1415–1420.

Lui JH, Hansen D V., Kriegstein AR. 2011. Development and evolution of the human neocortex. Cell.

Martín C, Bueno D, Alonso MI, Moro JA, Callejo S, Parada C, Martín P, Carnicero E, Gato A. 2006. FGF2 plays a key role in embryonic cerebrospinal fluid trophic properties over chick embryo neuroepithelial stem cells. Dev Biol. 297:402–416.

Oliphant TE. 2007. Python for scientific computing. Comput Sci Eng. 9:10–20.

Pedregosa F, Varoquaux G, Gramfort A, Michel V, Thirion B, Grisel O, Blondel M, Prettenhofer P, Weiss R, Dubourg V, Vanderplas J, Passos A, Cournapeau D, Brucher M, Perrot M, Duchesnay É. 2011. Scikit-learn: Machine Learning in Python. J Mach Learn Res. 12:2825–2830.

Rajagopalan V, Scott J, Habas PA, Kim K, Corbett-Detig J, Rousseau F, Barkovich AJ, Glenn OA, Studholme C. 2011. Local tissue growth patterns underlying normal fetal human brain gyrification quantified in utero. J Neurosci. 31:2878–2887.

Rajagopalan V, Scott J, Habas PA, Kim K, Rousseau F, Glenn OA, Barkovich AJ, Studholme C. 2012. Mapping directionality specific volume changes using tensor based morphometry: an application to the study of gyrogenesis and lateralization of the human fetal brain. Neuroimage. 63:947–958.

Reillo I, de Juan Romero C, García-Cabezas MÁ, Borrell V. 2011. A role for intermediate radial glia in the tangential expansion of the mammalian cerebral cortex. Cereb Cortex. 21:1674–1694.

Sarnat HB, Flores-Sarnat L. 2016. Telencephalic Flexure and Malformations of the Lateral Cerebral (Sylvian) Fissure. In: Pediatric Neurology. Elsevier Inc. p. 23–38.

Tallinen T, Biggins JS. 2015. Mechanics of invagination and folding: Hybridized instabilities when one soft tissue grows on another. Phys Rev E - Stat Nonlinear, Soft Matter Phys. 92.

Tallinen T, Chung JY, Biggins JS, Mahadevan L. 2014. Gyrification from constrained cortical expansion. Proc Natl Acad Sci U S A. 111:12667–12672.

Tallinen T, Chung JY, Rousseau F, Girard N, Lefèvre J, Mahadevan L. 2016. On the growth and form of cortical convolutions. Nat Phys. 12:588–593.

Thompson PM, Schwartz C, Lin RT, Khan AA, Toga AW. 1996. Three-dimensional statistical analysis of sulcal variability in the human brain. J Neurosci. 16:4261–4274.

Tustison NJ, Avants BB. 2013. Explicit B-spline regularization in diffeomorphic image registration. Front Neuroinform. 7:39.

Van D.C. E. 1997. A tension-based theory of morphogenesis and compact wiring in the central nervous system. Nature. 385:313–318.

Walsh CA. 1999. Genetic malformations of the human cerebral cortex. Neuron.

Welker W. 1990. Why Does Cerebral Cortex Fissure and Fold? p. 3–136.

West J, Newton PK. 2019. Cellular interactions constrain tumor growth. Proc Natl Acad Sci U S A. 116:1918–1923.

Wright R, Kyriakopoulou V, Ledig C, Rutherford MA, Hajnal J V., Rueckert D, Aljabar P. 2014. Automatic quantification of normal cortical folding patterns from fetal brain MRI. Neuroimage. 91:21–32.

Xu G, Knutsen AK, Dikranian K, Kroenke CD, Bayly P V., Taber LA. 2010. Axons pull on the brain, but tension does not drive cortical folding. J Biomech Eng. 132.

Yushkevich PA, Piven J, Hazlett HC, Smith RG, Ho S, Gee JC, Gerig G. 2006. User-guided 3D active contour segmentation of anatomical structures: significantly improved efficiency and reliability. Neuroimage. 31:1116–1128.

Zilles K, Palomero-Gallagher N, Amunts K. 2013. Development of cortical folding during evolution and ontogeny. Trends Neurosci.

